# Compressed inverted indexes for scalable sequence similarity

**DOI:** 10.1101/2025.11.21.689685

**Authors:** Florian Ingels, Léa Vandamme, Mathilde Girard, Clément Agret, Bastien Cazaux, Antoine Limasset

## Abstract

Modern sequencing continues to drive explosive growth of nucleotide sequence archives, pushing MinHash sketching methods to their scalability limits. Tools such as Mash, Dashing2, and Bindash2 provide compact sketches and accurate similarity estimates, but they typically use forward indexes that materialize each sketch as an explicit fingerprint vector. This architecture makes large-scale similarity search and collection-versus-collection comparison costly in both time and memory, especially at the scale of millions of sequences.

We revisit sketch index architectures and develop a framework based on inverted indexes over sketch fingerprints. We introduce a cost model for sketch comparison and prove that, with suitably compressed posting lists, inverted indexes can match the asymptotic space complexity of forward indexes. Using this model, we design all-vs-all comparison algorithms between two inverted indexes whose running time is proportional to the total number of matching sketch positions, yielding output-sensitive optimality and enabling efficient large collection comparisons at scale.

Because many applications impose similarity thresholds, we add two early-pruning schemes for Jaccard similarity. The first is exact and eliminates pairs guaranteed not to reach a target threshold. The second is probabilistic and exploits partial match statistics to discard pairs unlikely to exceed the threshold, with explicit control of the false-rejection probability. These schemes reduce time and memory while preserving rigorous guarantees on retained high-similarity pairs.

We implement these ideas in Onika, an open-source Rust system built on compressed inverted posting lists at github.com/Malfoy/Onika. Onika also applies similarity-aware document reordering to shrink index size and improve locality, especially for redundant collections. Experiments on bacterial genome repositories and long-read HiFi datasets show that Onika matches or improves sketch sizes of leading tools while accelerating large-scale search and collection-versus-collection comparison by up to several orders of magnitude in low-redundancy regimes, without compromising sensitivity at practically relevant similarity thresholds.

## 1 Introduction

One of the foremost challenges in modern computational biology is the large-scale analysis of sequence data. The exponential growth of genomic databases, a trend observed for decades, has accelerated dramatically with the advent of high-throughput sequencing. Public repositories like the Sequence Read Archive (SRA) now house hundreds of petabases of raw data, and the number of assembled genomes is in the hundreds of millions [1, 2]. This data deluge shows no signs of abating [3]. Continuous improvements in sequencing technology are making data generation cheaper, faster, and more accessible, with portable sequencers enabling real-time genomics in diverse environments [4]. Furthermore, future technologies such as Sequencing by Expansion [5] and 3D sequencing promise to increase data output by orders of magnitude, with projected throughputs nearing gigabases per second [6, 7]. This wealth of information represents a goldmine for biological discovery, powering fields like pangenomics, large-scale phylogenetics, and metagenomic characterization, but only if it can be effectively queried and analyzed.

The sheer volume of this data renders traditional, alignment-based comparison methods like BLAST computationally prohibitive. To manage this scale, the field has widely adopted alignment-free approaches based on *k*-mers, which are short substrings of length *k* [8]. The similarity between two sequences can be efficiently approximated by comparing their sets of *k*-mers, typically using the Jaccard index [9]. This metric serves as a robust proxy for established biological measures like Average Nucleotide Identity (ANI), a cornerstone of microbial taxonomy [10]. However, comparing the complete *k*-mer sets of millions of genomes remains an intractable task [11]. To circumvent this, an intermediate generation of methods represented entire *k*-mer sets using probabilistic data structures [12]. Approaches relying on Bloom filters [13, 14, 15], XOR filters [16], or Minimal Perfect Hash Functions [17, 18, 19] (MPHFs) allowed for compressed, queryable representations of a sequence’s full *k*-mer content. These structures traded a controllable false-positive rate for significant memory savings, but an even more aggressive form of compression was needed for the next leap in scale.

This next step was to abandon the representation of the full *k*-mer set and instead rely on a small, selective subsample, a technique now known as sketching. The seminal MinHash algorithm demonstrated that the Jaccard index could be accurately estimated from these compact, fixed-size “sketches” [20]. This breakthrough spurred the creation of a generation of powerful genomic analysis tools. Mash was the first to popularize MinHash for rapid, large-scale genome distance estimation [9]. Subsequent tools have continuously refined this core concept. Bindash [21, 22], for example, introduced innovations like using partition-MinHash with optimal densification and the use of smaller, *b*-bit fingerprints to enhance memory efficiency. Dashing [23, 24] explored the use of HyperLogLog fingerprints, and more recently, Dashing2 employed SetSketches [25] to incorporate *k*-mer multiplicities. It is important to note that these tools all produce fixed-size sketches, whose memory footprint is independent of the input genome’s size. A distinct lineage of subsampling techniques instead builds scalable sketches that grow with the input sequence, either by selecting *k*-mers through hash-threshold rules such as modulo-MinHash and FracMinHash [26, 27, 28], which underlie scaled signatures in sourmash and are also leveraged in skani [10], or by performing selection at the minimizer level [29]. These scalable sketches are particularly useful when the relevant similarity is asymmetric and one wishes to approximate containment rather than Jaccard similarity, as in metagenomic screening scenarios [30], however our focus here is on the fixed-size sketching paradigm, which has become dominant for rapid all-vs-all database comparison [31].

Despite their distinct innovations in fingerprinting strategies and statistical accuracy, all these fixedsize sketching methods are fundamentally concatenations of per-document sketches. Consequently, to identify documents similar to a given query, its sketch must be compared against every sketch in the index. Because each sketch has the same size, the sketch-to-document mapping is implicit; we therefore refer to such index structures as forward indexes. Although each individual comparison involves inexpensive integer operations that can be heavily optimized using SIMD instructions [22], the overall query time still scales linearly with the number of database entries. This computational complexity, expressed as *O*(*N* · *S*) where *N* is the number of queried sequences and *S* is the sketch size, represents a critical scalability bottleneck. As genomic databases grow from millions to potentially billions of entries, an approach whose cost grows linearly with database size is fundamentally unsustainable. Even worse, comparing a collection of *Q* sequences versus another collection of *R* sequences leads to a quadratic query time in *O*(*Q* · *R* · *S*), thus making this kind of global comparison intractable with large databases.

In our recent work, we introduced NIQKI, a novel indexing framework designed to dismantle this scalability barrier [11]. The key innovation of NIQKI is its departure from the forward index in favor of an inverted index [32]. Analogous to the index in a textbook, which maps terms to the pages they appear on, our inverted index maps each possible fingerprint value to a list of all documents that contain it. When a query is performed, NIQKI retrieves only the lists corresponding to the fingerprints present in the query sketch and aggregates the matches. Instead of being linear in the size of the database, the computational cost of a given query becomes proportional to the number of fingerprint matches, which is optimal in the sense that the necessary work is done (i.e. finding all matches) but no additional work is done for “nothing”. This is directly linked to the concept of “output polynomial time” used to evaluate the complexity of enumeration algorithms [33]; in our case, with output linear time.

However, this proof of concept approach, while being orders of magnitude faster than state of the art, showed exponential memory overhead and potentially expensive memory representation. In this paper we analyze the expected memory cost of inverted indexes and show that we can expect such an index to have the same memory cost as a forward index while conserving arguably optimal query times showing that they can be optimal both in time and memory. We present an efficient implementation of such a strategy in a tool Onika implemented in idiomatic Rust and show that it is able to outperform the state of the art in both time and memory at scale. Finally, we explore the importance of ordering documents to further optimize the performance of such indexes.

**Remark 1**. *It should be noted that although inverted indexes are not really exploited in bioinformatics (mainly because of the perceived memory penalty associated with them, which we refute in this article), they are already being successfully exploited in other disciplines with similar applications. In particular, the so-called “set-similarity join” problem has been widely studied in the data mining, information retrieval and database communities, especially in the context of web document clustering. To determine pairs of sets with high enough similarity (Jaccard or cosine), inverted indexes have been successfully used for more than 20 years — see for example [34, 35, 36*].

## 2 Methods

### 2.1 Forward index analysis

#### 2.1.1 Sketches

Let *S* and *W* be two integers. A *sketch s* is a tuple of *S* fingerprints, where a fingerprint is a *W*-bit integer, i.e. *s* = (*f* ^(1)^, …, *f* ^(*S*)^) with 0 ≤ *f* ^(*i*)^ ≤ 2^*W*^ − 1. Sketches are used to represent a large set of elements, and have been introduced to estimate Jaccard indices between sets [20]. The main interest of sketches is that they allow for very quick comparisons since they are orders of magnitude smaller than the set they actually represent. Three main approaches have been introduced to compute a sketch from a set of *U* elements [37]:

##### Multiple hash functions

Suppose we have *S* distinct hash functions *h*_1_, …, *h*_*S*_. For 1 ≤ *i* ≤ *S, f* ^(*i*)^ is defined as the smallest hash value, using *h*_*i*_, among all elements of the set. Computing the sketch requires *O*(*US*) time and sketch comparison is *O*(*S*).

##### Several minimal values

Using only one hash function, the sketch is defined as the *S* smallest values among the hashed elements. Sketch construction becomes *O*(*U* log *S*).

##### Partitions

Again, the elements are hashed using a single hash function. The hash values are partitioned into *S* buckets (e.g. by their first bits), and the sketch is made of the smallest elements of each partition. Construction is reduced to *O*(*U*) and comparison to *O*(*S*).

In this work, we resort to the last option (see Section 2.3) and refer interchangeably to sketches or partitions in the sequel. Literature offers many options for constructing and choosing the fingerprints (*b*-bit MinHash [21], HyperLogLog [23], HyperMinHash [11], SetSketch [24], etc.). At this stage, we do not assume how they are constructed; later on, we will require fingerprints to satisfy a uniformity condition, which will be discussed when introduced.

#### 2.1.2 Comparing forward indexes

When dealing with a set 𝒟 of distinct documents (i.e. sets of elements), and regardless of the construction of the sketches for each of these documents, the standard practice is to build a forward index (using Knuth’s terminology [38]), that is, an array of dimensions *D* × *S* — with *D* = |𝒟| — where the row of index *i* contains *s*_*i*_, the sketch of the *i*-th document. We denote by 𝒟_*F*_ the forward index built from 𝒟. Since each sketch contains *S* values encoded with *W* bits, the size of 𝒟_*F*_ is *O*(*DSW*) bits.

Provided two documents represented by their sketches *q* and *r*, the Jaccard index between these documents can be estimated from the value 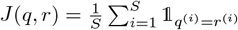 where *q*^(*i*)^ (resp. *r*^(*i*)^) represents the *i*-th fingerprint of sketch *q* (resp. *r*). The task usually performed in practice is to compare all pairs of documents coming from two sets of documents 𝒬 and ℛ (of respective cardinality *Q* and *R*) — sometimes with 𝒬 = ℛ. Therefore, the goal is to compute a similarity matrix *M* of size *Q* × *R* where *M* [*q, r*] = *S* · *J* (*q, r*) — *i*.*e*. entry (*q, r*) of the matrix counts the number of matches between the sketches *q* and *r*. The Jaccard index is then straightforwardly estimated by dividing the matrix by *S*.

Using forward indexes, the only way to compute such matrix *M* is to parse the entire indexes 𝒬_*F*_ and ℛ_*F*_ and compare each sketch of 𝒬 against each sketch of ℛ. This can be accomplished using Algorithm 1. Examples of forward indexes and their comparison can be found in Figures 1a and 1c, respectively. To analyze the cost of computing *M* using Algorithm 1, we introduce the total score Σ_*M*_ of the matrix, which is Σ_*M*_ = Σ_*q*∈𝒬_ Σ_*r*∈ℛ_ *M* [*q, r*]. We also introduce the following cost model.

**Figure 1:**
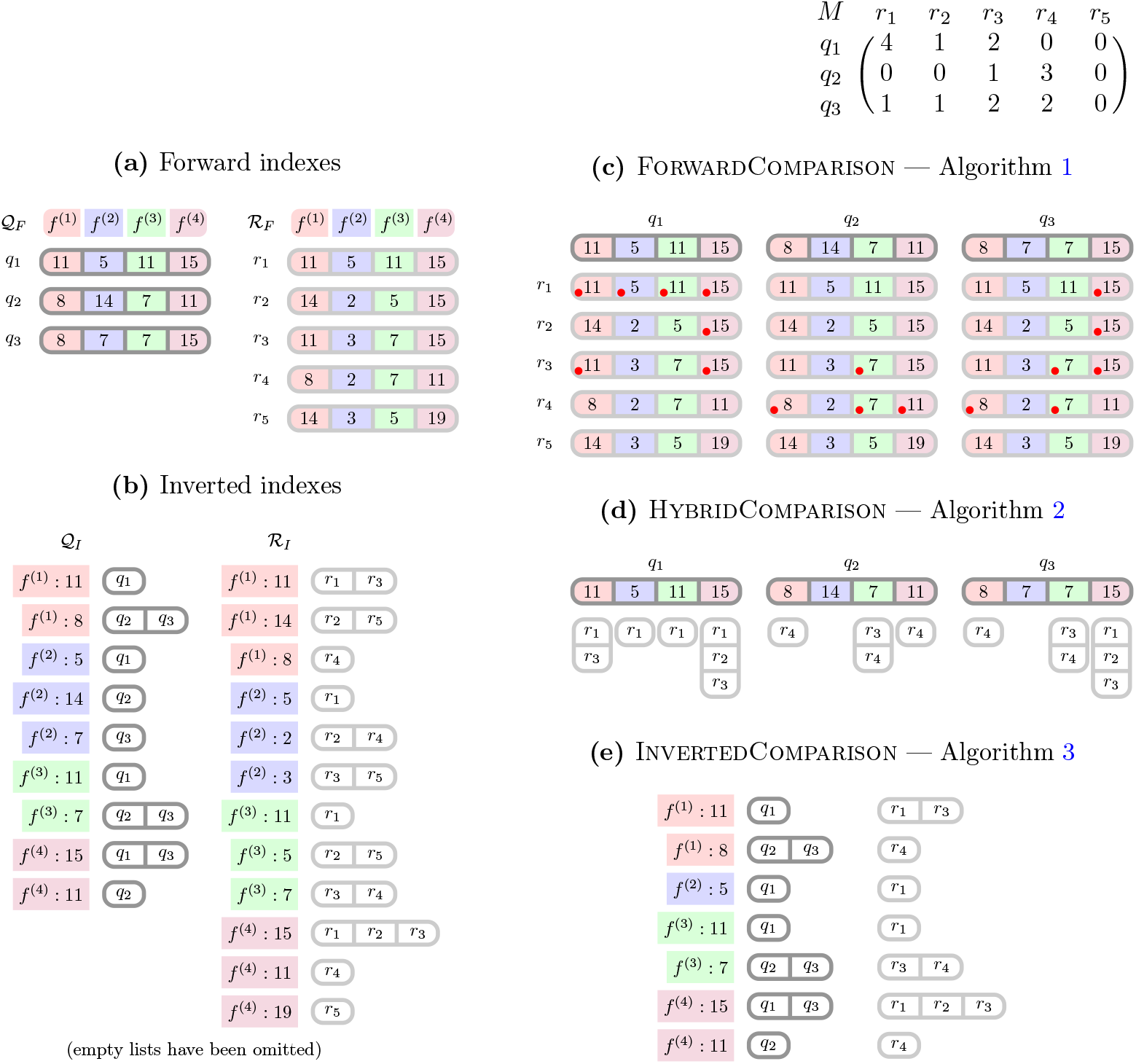
Examples of indexes and comparisons of two sets of documents 𝒬 and ℛ, with *Q* = 3, *R* = 5, and *S* = 4. In (c), red dots indicate a match. The comparison matrix *M*, with a total score Σ_*M*_ = 17, is given on the right.

##### Definition 1

(Cost model). *Let us denote by c*_*c*_ *the machine cost of a comparison, by c*_*i*_ *the machine cost of an increment, and by c*_*a*_ *the machine cost of a random access. We suppose that the machine cost of sequential access is negligible*.

We provide in Appendix S2 an alternative model that takes into account the machine cost of sequential access, as well as the complexities for upcoming Algorithms 1, 2 and 3 under this new model — adding this cost does not change any conclusion we draw in the sequel about their compared performances. For now, we have the following result.

##### Proposition 1

*Using our cost model, Algorithm 1 runs in QRS* · *c*_*c*_ + Σ_*M*_ · *c*_*i*_ *time*.

*Proof*. Obviously, as per the pseudocode, we need to perform *QRS* comparisons; however we perform in an increment if and only if a match is found, the number of which is exactly the sum of the entries of the matrix *M*.

### 2.2 Inverted index analysis

#### 2.2.1 Inverted index

##### Algorithm 1

ForwardComparison

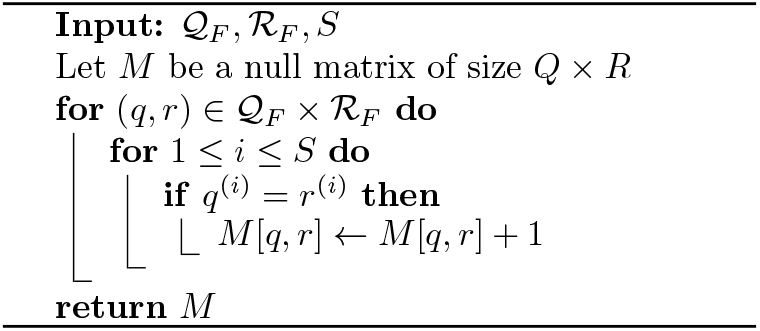

Consider a set of documents 𝒟 (with *D* = |𝒟|) and their sketches. The inverted index (again, using Knuth’s terminology [38]) 𝒟_*I*_ built from 𝒟 can be seen as a tuple of *S* separate indexes, 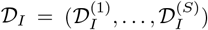, where, for 1 ≤ *i* ≤ *S*, the subindex 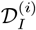 is an array of size 2^*W*^ where cell 0 ≤ *f* ≤ 2^*W*^ – 1 contains the indices of the documents of 𝒟 whose *i*-th fingerprint is *f*. In other words,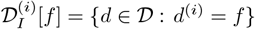. Note that this list is potentially empty. We investigate the actual construction of an inverted index in Section 2.3.

The inverted index contains *S* × 2^*W*^ lists of indices of documents, of varying lengths. Let us denote size 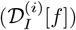 the size, in bits, needed to encode the list 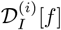. Without loss of generality, 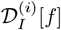 can be represented as a sorted list of indices. Equivalently, we can encode this list using *δ*-encoding, that is registering (in binary) the first index as is, and the subsequent indices as their difference from the previous index. Using such representation, we have the following result.

##### Theorem 1

*For any* 1 ≤ *i* ≤ *S, assuming all fingerprints have an equal chance of being chosen to represent a given document, that is, for all* 0 ≤ *f* ≤ 2^*W*^ − 1 *and d* ∈ 𝒟, 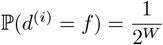, *and provided that D is large enough, we have* 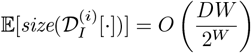 *bits using δ-encoding*.

*Proof*. The proof is deferred to Appendix S1.

Assuming fingerprint uniformity is a strong and restrictive requirement for inverted-indexing sketches. In particular, it is violated by several widely used state-of-the-art sketches, including HyperLogLog and related mergeable cardinality sketches, as well as HyperMinHash and SetSketch, whose fingerprints are inherently non-uniform and often highly compressible [39], making them poor candidates for inverted-index schemes. In contrast, *b*-bit MinHash fingerprints, obtained by keeping the *b* least significant bits of large (approximately uniform) hash values, can be treated as uniform under standard assumptions [40]. Moreover, the optimal fingerprint size depends on the expected Jaccard similarity; for highly similar documents (Jaccard *>* 0.5), the optimal size can be as small as a single bit [41], which justifies using small, fixed-width fingerprints in practice and explains why this fingerprint is implemented in both Dashing2 and Bindash.

Since the expected size of a list is 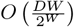 by Theorem 1 and since there are *S*2^*W*^ such lists, we get that the inverted index can be stored in *O*(*DSW*) bits, which is identical to the size of the forward index as established before. In other words, by selecting appropriate fingerprints and using *δ*-encoding, we show that we can get rid of the overhead traditionally associated with inverted indexes (and found in our previous tool Niqki). Examples of inverted indexes are provided in Figure 1b.

#### 2.2.2 Comparing inverted indexes

Let 𝒬 and ℛ be two sets of documents of cardinality, respectively, *Q* and *R* — as previously. We are interested in computing the similarity matrix *M* as before. Let ℛ_*I*_ (resp. 𝒬_*I*_) be the inverted index built from ℛ (resp. 𝒬). A first option to compute *M* is to use a hybrid approach, using the forward index 𝒬_*F*_ and the inverted index ℛ_*I*_, as was done in Niqki. Algorithm 2 describes this approach, with Proposition 2 providing its temporal complexity and Figure 1d an example.

##### Proposition 2

*Using our cost model, Algorithm 2 runs in QS* · *c*_*a*_ + Σ_*M*_ · *c*_*i*_ *time*.

*Proof*. For each fingerprint *q*^(*i*)^ of 𝒬_*F*_, we need to perform a random access to obtain the list 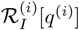 — there are *QS* of them. Now, since 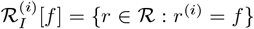, notice that equivalently 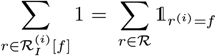. Therefore, when considering the cost of increment, we have — using interchangeably 𝒬 and 𝒬_*F*_ as an abuse of language:

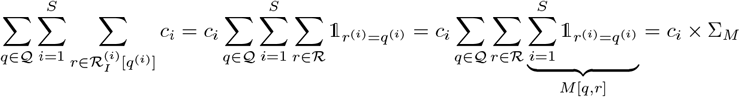

##### Algorithm 2

HybridComparison

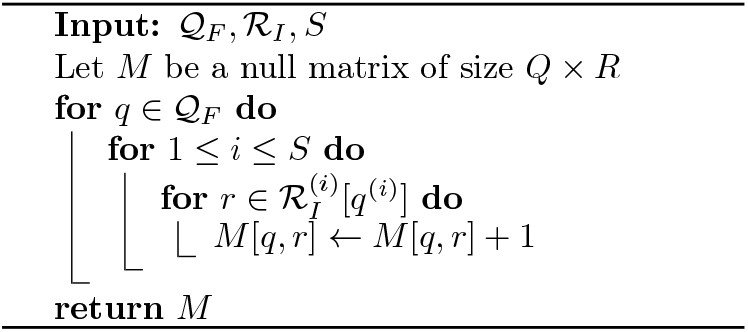

##### Algorithm 3

InvertedComparison

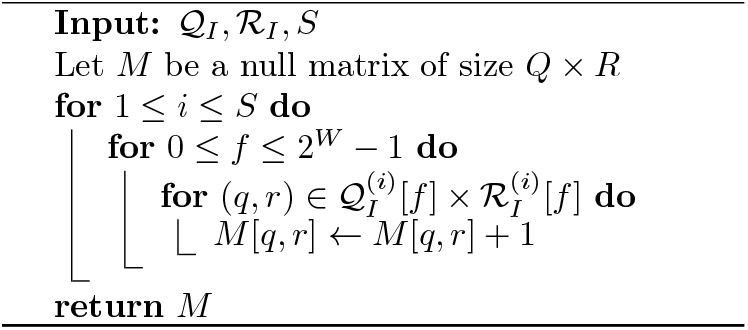

Finally, if the inverted index for 𝒬 has been built, that is, 𝒬_*I*_, then there exists an approach that is optimal, described in Algorithm 3 and whose complexity is given in the following Proposition. Figure 1e provides an example.

##### Proposition 3

*Using our cost model, Algorithm 3 runs in* Σ_*M*_ · *c*_*i*_ *time*.

*Proof*. Notice that no random accesses are performed, so we only need to account for the cost of increments. We use the same trick as of the proof of Proposition 2, that is, for any set of documents 𝒟 and any fingerprint *f*,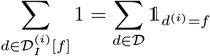. Then, we have

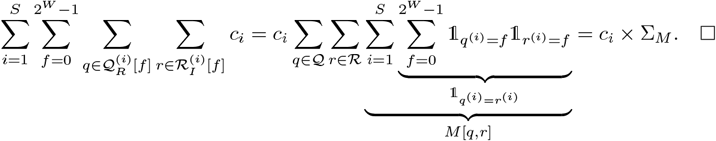

A comparative table of the different algorithms and their complexities can be found in Table 1. We can see that Algorithm 3 is the best of the three approaches considered. Furthermore, since the goal of all three algorithms is to find all matches — i.e., Σ_*M*_ — Algorithm 3 is actually optimal in the sense that it only computes what is requested and does not perform any unnecessary extra work. Note that even under the alternative cost model available in Appendix S2, taking into account the cost of sequential access, the same conclusion holds.

**Table 1:** Summary of the complexities of the comparison algorithms under our cost model.

|  |  |  |  |
| --- | --- | --- | --- |
| Algorithm 1 (forward) | $QRS \cdot c_c + \Sigma_M \cdot c_i$ | (Proposition 1) | $O(QRS + \Sigma_M)$ |
| Algorithm 2 (hybrid) | $QS \cdot c_a + \Sigma_M \cdot c_i$ | (Proposition 2) | $O(QS + \Sigma_M)$ |
| Algorithm 3 (inverted) | $\Sigma_M \cdot c_i$ | (Proposition 3) | $O(\Sigma_M)$ |

In the end, from a theoretical point of view, using an inverted index is optimal both in terms of space (as it requires the same space as a forward index – from Theorem 1) and in terms of comparison time (from Table 1). Empirical evidence for this is presented in Section 3.

### 2.3 Sketching

We employ Nthash, an efficient rolling hash algorithm dedicated to efficient *k*-mer processing [42]. It is worth noting that Nthash’s flexibility would allow to construct sketches on spaced seeds as an alternative to contiguous *k*-mers [43]. A naive construction of the inverted index would allocate *S* · 2^*W*^ posting lists and fill them on the fly while computing the sketches. This induces severe memory fragmentation and a high peak memory footprint. Instead, we adopt a compact two-pass strategy.

In the first pass, we preallocate an array of size *S* · *N* (where *N* is the number of datasets) to store all fingerprints. While parsing each dataset, we compute its *S* fingerprints and store them in a transposed layout: rows correspond to partitions and columns to datasets. Once this array is filled, we scan it row by row in a second pass. Each row then provides, for a given partition, the *N* fingerprints together with their implicit dataset identifiers (the column indices). For each partition and each of the 2^*W*^ possible fingerprint values, we append dataset identifiers in increasing order, without any sorting step, since datasets are processed in a fixed global order. For a given partition, once all posting lists are built, they are *δ*-encoded and compressed to disk before processing the next partition. This reduces the working memory usage by a factor *S* compared to maintaining all *S* · 2^*W*^ lists simultaneously. On disk, the sketch representation for each partition consists of the compressed posting lists organized by fingerprint value. Since the number of distinct *k*-mers *U* is typically much larger than the sketch size *S* (i.e., *U* » *S*), the cost of sketch computation is usually dominated by input parsing and I/O rather than hashing. As discussed earlier, we expect these inverted sketches not to exceed the size of forward sketches that store fingerprints explicitly as integers. In settings with substantial redundancy between datasets, compression can be further improved by reordering dataset identifiers so that similar datasets receive nearby identifiers. This increases locality in the posting lists and improves *δ*-encoding, in the same spirit as the read index K2R [44], which reorders reads using Oreo [45] to reduce index size.

To exploit this effect, we implement an optional reordering step. We first estimate pairwise similarities between sketches using only a limited number of partitions (100 by default), which provides a coarse but inexpensive similarity approximation. We then construct an ordering greedily: starting from an arbitrary sketch, we iteratively choose as successor the most similar sketch not yet placed. The resulting permutation is applied to dataset identifiers before index construction, yielding more compressible posting lists at negligible additional cost.

#### 2.3.1 Comparison

To compare two collections, we first compute their respective sketches, which enables the following efficient procedure. The resulting inverted indexes are organized by partition and, within each partition, by fingerprint value. This layout allows both indexes to be scanned in a single sequential pass.

During this joint scan, for each partition and each fingerprint, if both collections contain a nonempty posting list, we update the similarity scores of all dataset pairs in the Cartesian product of the two corresponding identifier lists. The procedure is memory-efficient at the index level, since only the current posting lists from both collections need to be held in memory. Parallelism is achieved using a coarse-grained strategy, assigning disjoint subsets of partitions to different threads.

The main bottleneck is the score matrix itself, which requires *O*(*Q* · *R*) integer entries for *Q* queries and *R* reference datasets. For large collections, this matrix can dominate memory usage and induce frequent cache misses, as each increment may touch a distant memory location. This issue explains the high memory requirements observed in tools such as Dashing2 when scaling to large collections. Bindash2 mitigates this by processing datasets in chunks so that each partial score matrix fits into cache, but this design induces a quadratic number of chunk comparisons as *Q* and *R* grow, which leads to substantial runtime overhead.

We now aim to introduce a comparison strategy that is scalable in both time and memory. Our approach starts from the observation that sketch-based estimators are most effective for identifying highly similar pairs, where the Jaccard index is large, while their relative error increases as similarity decreases. In practice, sketches are predominantly used for tasks such as clustering or nearest-neighbor search, where only pairs above a user-defined similarity threshold are of interest and low-similarity pairs are ignored. We argue that this explicit similarity threshold can be exploited to avoid maintaining and updating a dense *O*(*Q* · *R*) score matrix and to prune unpromising candidates during the comparison process, yielding significant computational savings without affecting the set of relevant high-similarity pairs. The main idea is to replace the matrix *M* with a dictionary whose keys are the pairs (*q, r*) to be kept and whose values are *M* (*q, r*), thus reducing the memory usage to its strict minimum to achieve downstream applications. Let us denote by *t* ∈ [0, 1] this similarity threshold — *i*.*e*. we want to keep only pairs (*q, r*) for which *J* (*q, r*) ≥ *t*, or, equivalently, for which *M* [*q, r*] ≥ *tS*. The main challenge is therefore to cut-off pairs (*q, r*) that will not reach the threshold as quickly as possible — and thus reduce both the final memory footprint and the computation time.

First, there is a deterministic rule. Suppose we have encountered *k* matches in the first *n* partitions; if *k* +(*S* − *n*) < *tS*, then surely *M* [*q, r*] < *tS*, as there are not enough sketches left to consider to reach the required score. However, this rule requires *n* to be quite large to have an effective cut-off, where we would like to make the decision to reject the pair (*q, r*) as soon as possible. Therefore, we propose to resort to a probabilistic heuristic, at the risk of rejecting pairs (*q, r*) that would have passed the threshold.

Consider the vector of size *S* indicating, at position *i*, whether *q*^(*i*)^ = *r*^(*i*)^ or not. We can see this vector as *S* independent trials following a Bernoulli distribution of parameter *p* = *J* (*q, r*) (whose value is unknown), so that *M* (*q, r*) follows a binomial distribution of parameters (*S, p*). Let *X* be the random variable counting the number of matches encountered in the first *n* sketches; we have *X* ~ ℬ (*n, p*). Assuming that the pair (*p, q*) would indeed pass the threshold (i.e. *p* ≥ *t*), we want to estimate the probability ℙ (*X* = *k* | *p* ≥ *t*). If this probability is below another probabilistic threshold *s* ∈ [0, 1], then we reject the pair (*p, q*) as we estimate it implausible that the pair would in fact pass the threshold *t*. In practice, we do not use the actual probability ℙ (*X* = *k* | *p* ≥ *t*) but an upper bound, obtained as follows. Using the law of total probability, and assuming that *p* is a continuous random variable chosen at random for the pair (*q, r*), with (unknown) density *F*_*p*_, then

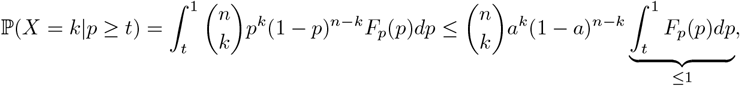

where *a* = max(*t, k/n*). Indeed, the function *p* ↦ *p*^*k*^(1 − *p*)^*n*−*k*^ is increasing on the interval [0, *k/n*[,maximal at *k/n*, and decreasing on]*k/n*, 1]. Therefore, if *t* ≤ *k/n*, its maximum is reached at *k/n*, and at *t* otherwise. To speed up the computation of 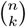, we instead use the recent and quite tight upper bound of Agievich [46]. For any 0 ≤ *k* ≤ *n*,

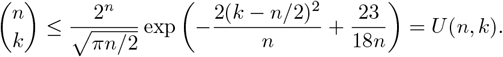

When *a* approaches 1, *U* (*n, k*)*a*^*k*^(1 − *a*)^*n*−*k*^ ≫ 1, therefore the heuristic will reject pairs (*p, q*) for which *k/n* and *t* are not too big, which is what we want in practice. In the end, we reject the pair (*p, q*) if and only if *U* (*n, k*)*a*^*k*^(1 − *a*)^*n*−*k*^ < *s*. Also, note that we can precompute the value *k*_*s*_ so that if *k* < *k*_*s*_, then *U* (*n, k*)*a*^*k*^(1 − *a*)^*n*−*k*^ < *s*. In this way, deciding whether or not to reject the pair (*q, r*) can be done in constant time. Appendix S3 provides some insights on the influence of the bound *U* (*n, k*), compared to 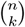, on the estimated probability.

A key point in our heuristic is to repeat this cut-off operation for different values of *n*. Once a pair (*p, q*) has been rejected, it is final; but if it passes the test, it is only temporary, until either (a) it is eventually rejected later due to insufficient matches, or (b) it reaches the required threshold *tS* and is definitively retained. Let us denote *n*_1_ < *n*_2_ < … the cut-off points, and 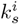 the required number of matches to pass the test at *n*_*i*_.

All three Algorithms 1, 2 and 3 involve a loop on 1 ≤ *i* ≤ *S*. While *i* ≤ *n*_1_, the algorithms run as written. For Algorithm 1, the adaptation is straightforward: when *i* = *n*_1_, if there has been 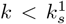 matches, we discard the pair and consider the next one, otherwise we resume the loop on *i* and perform the same operation at *n*_2_, …. For Algorithms 2 and 3, the adaptation is different. Instead of an actual matrix, pairs are stored as keys of a dictionary. When *i > n*_1_, next time we encounter a pair (*q, r*), one of the two following can occur:

- the pair (*q, r*) is not in the dictionary: either it has been discarded for failing the test, or it has never been seen (therefore obtaining 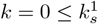 matches), and we ignore the pair (thus *M* [*q, r*] = 0);
- the pair (*q, r*) is in the dictionary with *k* matches. Let *n*_*j*_ be the largest cut-off point < *i*. If 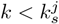, we discard the pair and delete the key from the dictionary. Otherwise, we increment *M* [*q, r*] to *k* + 1.

The effects in practice of this heuristic are discussed in upcoming Section 3.3.

## 3 Results

The experiments were performed on an Intel(R) Xeon(R) Gold 6430 system equipped with two 32-core sockets, 512 GB of RAM, and a 2 TB SSD.

### 3.1 Indexing large genome collections

As a first experiment, we evaluate the efficiency of Onika for large-scale whole-genome comparison, the canonical use case of sketch-based methods. We compare Onika against the best-performing tools Dashing2 and Bindash2. We exclude our previous implementation Niqki, which was designed for queryversus-collection scenarios, and Mash, which is outperformed by both Bindash2 and Dashing2 in recent benchmarks. In Figure 2 (left), we report sketching and comparison times on RefSeq bacterial genome collections of increasing size. Onika’s sketch construction is faster than both Bindash2 and Dashing2. More importantly, its comparison phase performs similarly on smaller collections and becomes up to 3-fold (Bindash2) and 5-fold (Dashing2) faster on the largest collections.

**Figure 2:**
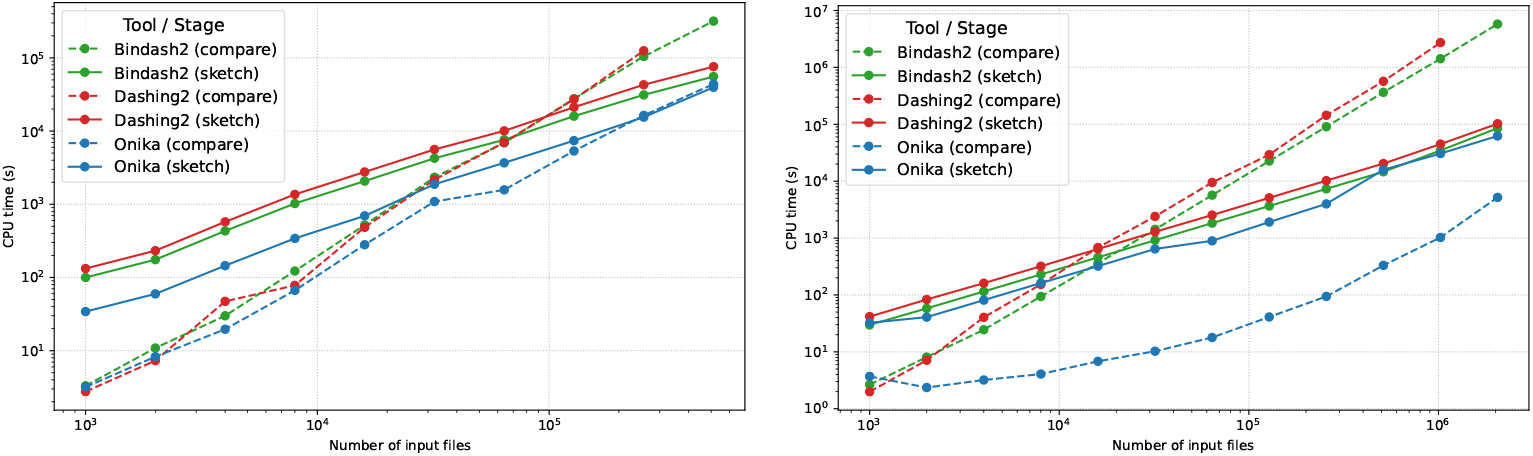
Comparison of CPU time of Dashing2, Bindash2 and Onika on a growing collection of bacterial genomes from RefSeq (left) and from random 1 Mb sequences (right). For both experiments *k* = 31 and *s* = 512.

However, this benchmark is pessimistic for Onika. Our approach scales with the total score Σ_*M*_ of the similarity matrix and is most penalized when many datasets are highly similar, since this inflates the number of candidate pairs that must be tracked. Bacterial collections are known to contain extensive redundancy and many near-duplicate genomes (90% of the ENA datasets were from only 20 genera, and 65% were from the five most common genera [13]), which makes this setting close to a worst-case scenario for Onika. To illustrate the opposite regime, Figure 2 (right) shows the same experiment on collections of independent random sequences of length 1 Mb. These datasets share no substantial similarity, which minimizes the total score Σ_*M*_ and exposes the best-case behavior of our strategy. In this setting, Onika is more than three orders of magnitude faster than the state of the art. While indexing random sequences is artificial, this experiment indicates the performance we can expect on future large, diverse, and nonredundant genome collections.

We previously argued that the inverted index representation used by Onika should not exceed the size of a forward index storing sketches explicitly. Figure 3 reports the sketch sizes for the RefSeq experiments above. To ensure a fair comparison, all sketches are compressed with `zstd −1`, which is Onika’s default. We observe that Onika’s sketch sizes are comparable to those of Bindash2, confirming the theoretical expectations in practice, and that the optional reordering step in Onika can reduce sketch size by more than 35%. In practice, we expect the reordering benefit to be highly dependent on the redundancy present in the collection. Memory usage results are presented in the Appendix (Figure 7) and show that Onika uses less memory than Dashing2 but is still more memory expensive than the almost constant memory usage of Bindash2 — we emphasize that this memory usage is directly linked to the size of all-vs-all matrices (remember that Bindash2 partition this matrix into small chunks, hence its limited memory usage) and not to the size of the indexes, that are adressed in Figure 3 (left) and Figure 4 (left).

**Figure 3:**
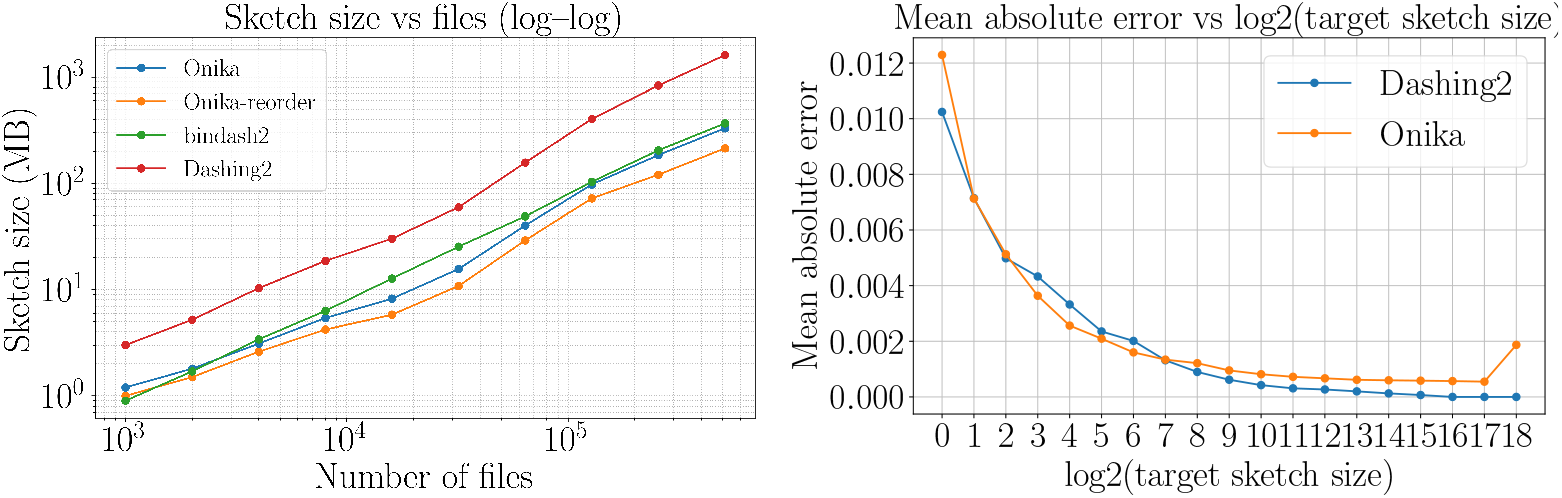
Comparison of sketch sizes of Dashing2, Bindash2 and Onika on a growing collection of RefSeq bacterial genomes (left). Qualitative comparison between Dashing2 and Onika versus a Jaccard ground truth according to the sketch size used on 2, 000 RefSeq bacterial genomes (right).

**Figure 4:**
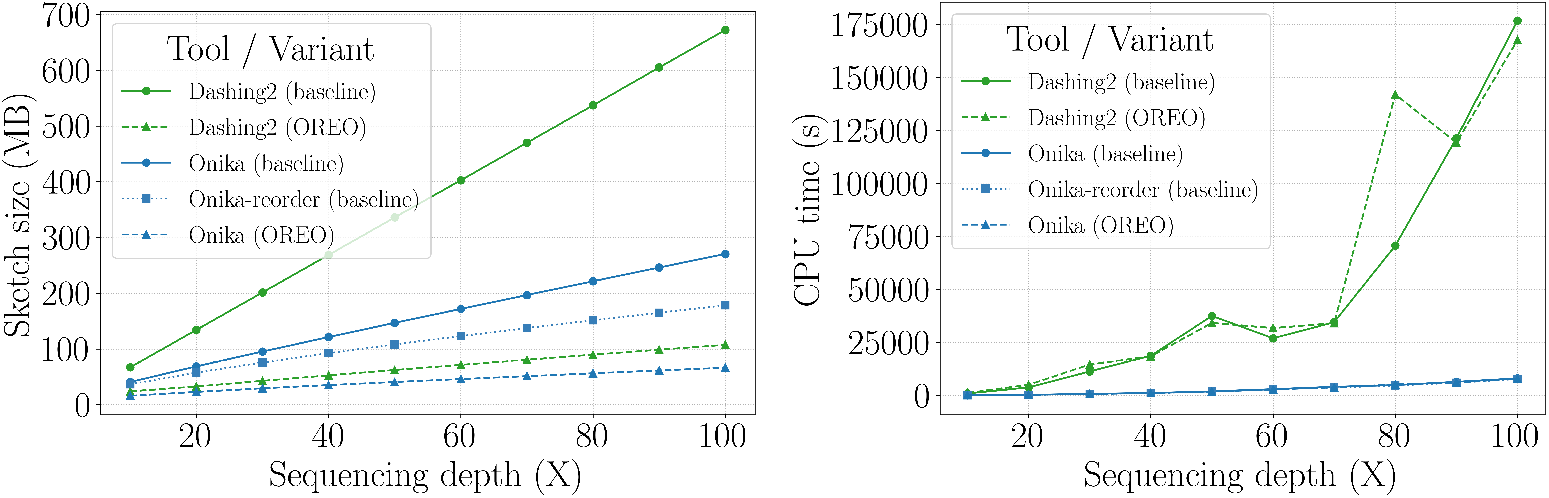
Comparison of sketch sizes (left) and CPU time (right) of Dashing2 and Onika, both with random original ordering and Oreo ordering, and finally Onika with its own reordering, for increasing depth of *A. thaliana* HiFi sequencing.

### 3.2 Indexing raw sequencing datasets

Another core application of sketching is the detection of similar reads within sequencing datasets for alignment-free overlap detection [47], assembly [48], error correction [49], and assignments [30]. We therefore evaluate Dashing2 and Onika on all-vs-all comparisons of HiFi read datasets. We excluded Bindash2 as it does not provide native support for this setting, and emulating it with a dedicated sketch per read would be computationally unrealistic. We also assess the impact of Onika’s optional reordering step on both sketch size and CPU time. Because read reordering is a standard technique for improving compression, we further apply Oreo [45], a tool that reorders reads to boost compressibility, and measure its effect on both methods. Figure 4 reports sketch sizes (Left) and CPU times (Right) for increasing coverage of *A. thaliana* HiFi data. For sketch size, Onika produces smaller sketches than Dashing2 under the original ordering, and its internal reordering further reduces size. When Oreo is applied, both tools benefit and converge to very similar sketch sizes, indicating that Onika can exploit either its own reordering or external compression-oriented orderings. For CPU time, Onika is consistently faster than Dashing2. Its performance is largely insensitive to read ordering, while Oreo dramatically degrades the running time of Dashing2 for reasons that remain unclear, highlighting an additional robustness advantage of Onika.

### 3.3 Heuristic efficiency

Finally, we evaluate the efficiency of the proposed heuristic pruning. In Figure 5 we report the number of detected hits for different similarity thresholds and probabilistic pruning levels, together with the corresponding Onika running times. Except under extremely aggressive probabilistic pruning (e.g. 0.1%), the fraction of missed hits remains negligible, while computation time decreases substantially. This confirms that the heuristic effectively accelerates comparisons without significantly affecting the set of relevant high-similarity pairs. In practice, the proportion of false negatives, hits that would have been reported without heuristic pruning, always remains below the chosen probabilistic threshold.

**Figure 5:**
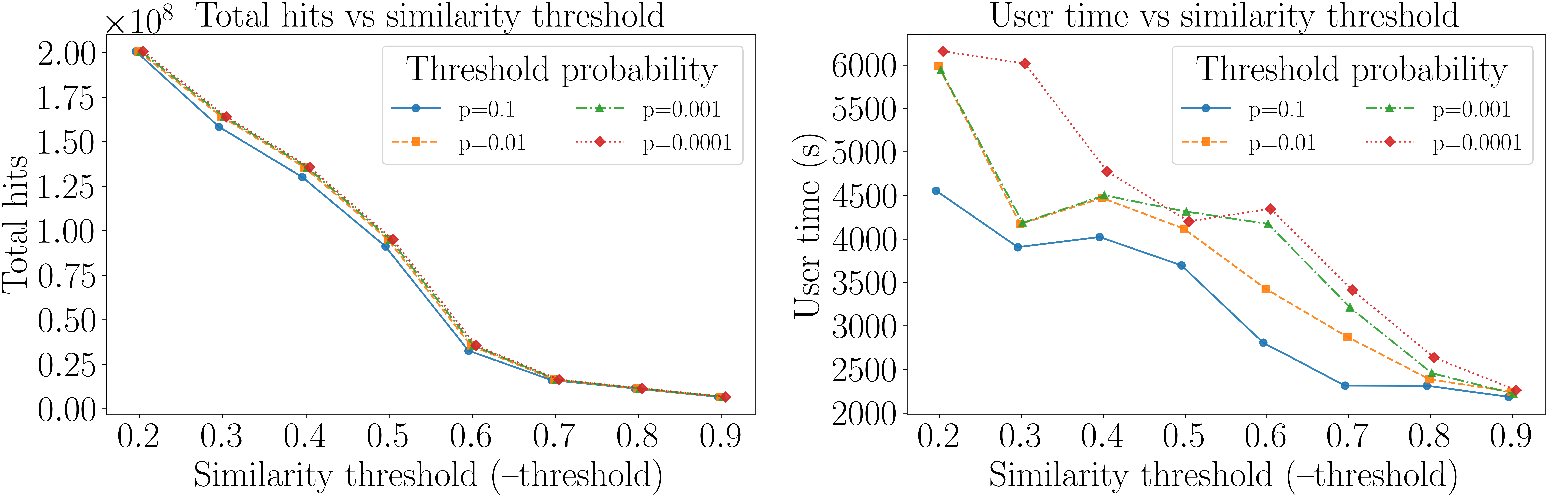
Evaluation of Onika’s heuristic pruning as a function of the similarity and probabilistic thresholds, reporting the number of hits (left) and runtime (right) on 100k RefSeq bacterial genomes. *P* = 0 denotes that no heuristic is used.

## 4 Conclusion

In this paper, we tackled the problem of making MinHash-style sketch comparison optimal at scale by recasting it as an inverted-index computation. We proved that inverted and forward sketches have identical storage requirements, *O*(*DSW*) bits (Theorem 1). We then introduced and analyzed a comparison cost model and we proved that all-pairs comparison achieved the irreducible Σ_*M*_ *c*_*i*_ cost when both sides were inverted — strictly improving over the forward and hybrid baselines *QRS c*_*c*_ + Σ_*M*_ *c*_*i*_ and *QS c*_*a*_ + Σ_*M*_ *c*_*i*_ (Propositions 1, 2 and 3).

We presented Onika, a Rust implementation that realizes these guarantees end to end. We designed a resource-efficient two-pass construction that built *δ*-compressed postings partition by partition, avoiding fragmentation and tightly bounding working memory, and we implemented a high-throughput inverted–inverted comparator that delivered the theoretical time bound in practice. We also introduced a unique, optional dataset-reordering pass that increases locality and amplifies delta compressibility, thereby shrinking sketch size without sacrificing throughput.

We developed a constant-time, threshold-aware pruning test: given a user-specified similarity threshold *t*, after processing *n* partitions, we rejected pairs with *k* < *k*_*s*_ matches, where *k*_*s*_ is computed so that an accepted pair (*i*.*e*. with Jaccard ≥ *t*) would obtain only *k* < *k*_*s*_ matches with probability < *s*. This rule preserved high-similarity pairs while sharply reducing candidate maintenance and update traffic.

We demonstrated empirically that Onika was competitive and scaled favorably on redundancy-heavy RefSeq bacterial genomes; that on diverse synthetic collections, where the effective score mass Σ_*M*_ was intrinsically small, it achieved orders-of-magnitude speedups; and that on read collections it outperformed Dashing2 in runtime while producing comparable or smaller sketches. Reordering steps further compressed sketches, with gains increasing alongside redundancy.

Natural directions for extension are stronger pruning with explicit error control; redundancy-aware mutualized computations to avoid repeated work across similar documents; algorithms specialized for top-*K* retrieval rather than dense distance matrices [50]; overlap-centric accelerations for read–read detection under tight neighborhood assumptions [48]; and GPU implementation [51] encompassing both partition-wise construction and merge-style inverted comparison.

## Acknowledgements

This work was supported by the French National Research Agency AGATE [ANR-21-CE45-0012] and full-RNA [ANR-22-CE45-0007]. With financial support from ITMO Cancer of Aviesan within the framework of the 2021-2030 Cancer Control Strategy, on funds administered by Inserm. The authors are grateful to three anonymous reviewers for their valuable comments on a first version of the manuscript.

## Supplementary material

### S1 Proof of Theorem 1

Without loss of generality, we assume *S* = 1 — so that we consider a single sketch per document *d* ∈ 𝒟, and we fix the fingerprint 0 ≤ *f* ≤ 2^*W*^ − 1. We are interested in compute the size, in bits, of the list {*d* ∈ 𝒟 : sketch(*d*) = *f* }, that we denote size(𝒟_*I*_ [*f*]) (dropping the subscript depending on *S*).

Let *B*_1_, …, *B*_*D*_ be a sequence of independent Bernoulli trials of parameter 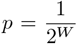. For *i* ≥ 1, let *T*_0_ = 0 and *T*_*i*_ = inf{*n > T*_*i*−1_ : *B*_*n*_ = 1} be the time at which the *i*-th success is found in the sequence; and let *G*_*i*_ = *T*_*i*_ − *T*_*i*−1_ be the number of trials between the (*i* − 1)-st success and the *i*-th success. The variables *G*_*i*_’s are independent, since the *B*_*i*_’s are. Let us denote 0 ≤ *τ* ≤ *D* the total number of successes, that i s, *τ* = inf{*n* ∈ ℕ : *T*_*n*_ *> D*}, assuming we can continue the sequence *B*_1_, *B*_2_, … beyond *D*. Since 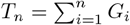, we conclude that *τ* is a stopping time with respect to the *σ*-algebra generated by the *G*_*i*_’s.

In our context, the variable *B*_*d*_ represents whether or not sketch(*d*) = *f*; the variables *T*_1_, …, *T*_*τ*_ correspond to the indices of the documents containing *f*, the variables *G*_1_, …, *G*_*τ*_ to the distance between those indices, and *τ* the number of documents that contain *f*. Therefore, if we encode the list of documents containing *f* using *δ*-encoding, the number of bits size (𝒟_*I*_ [*f*]) we need is expressed as the following random variable

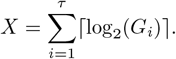

Since *τ* is a stopping time for the *G*_*i*_’s, it is also a stopping time for the ⌈log_2_(*G*_*i*_) ⌉’s; and noticing that 𝔼 [⌈log_2_(*G*_*i*_) ⌉] < ∞, using Wald’s equation [52], we get that

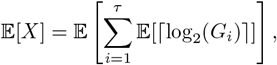

which is 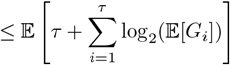, by Jensen’s inequality and concavity of log.

The exact distribution of the *G*_*i*_’s is established in [53], where it is also proven that the *G*_*i*_’s converge in distribution when *D* → ∞, to a geometric distribution of parameter *p*. Therefore, 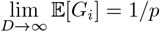 by denoting *C*_*D*_ = max_1≤*i*≤*D*_ 𝔼 [*G*_*i*_], we have 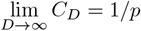.

Going back to *X*, we have 𝔼 [*X*] ≤ 𝔼 · [*τ*] (1 + log_2_(*C*_*D*_)). Obviously, *τ* follows a binomial distribution of parameters (*D, p*), so that 𝔼 [*X*] ≤ *Dp*(1 + log_2_(*C*_*D*_)). Finally, for *D* sufficiently large, there exists a constant *C >* 0 so that 1 + log_2_(*C*_*D*_) ≤ *C* log_2_(1*/p*), hence the result.

### S2 Runtime of Algorithms 1, 2 and 3 when taking into account the cost of sequential access

In this section, we investigate the complexities of Algorithms 1, 2 and 3 using our cost model — see Definition 1, but this time without assuming the machine cost of a sequential access is negligible; let us denote this cost by *c*_*s*_.

#### Lemma 1

*Algorithm 1 runs in O*(*QRS*) · *c*_*s*_ + *QRS* · *c*_*c*_ + Σ_*M*_ · *c*_*i*_ *time*.

*Proof*. We need to evaluate the following sum.

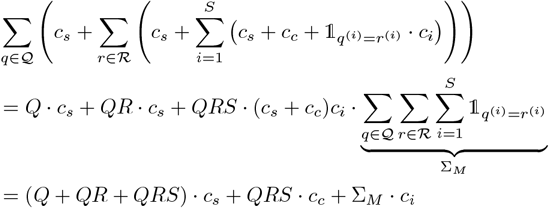

#### Lemma 2

*Algorithm 2 runs in O*(*QS* + Σ_*M*_) · *c*_*s*_ + *QS* · *c*_*a*_ + Σ_*M*_ · *c*_*i*_ *time*.

*Proof*. We need to evaluate the following sum

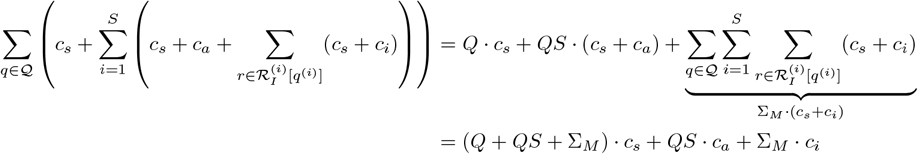

using the trick 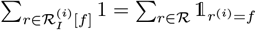.

#### Lemma 3

*Under the same assumption as Theorem 1 — that is, all fingerprints have an equal chance of being chosen to represent a given document; and provided that both Q and R are large enough, Algorithm 3 runs in O*(*S* + Σ_*M*_) · *c*_*s*_ + Σ_*M*_ · *c*_*i*_ *time*.

*Proof*. We need to evaluate the following sum:

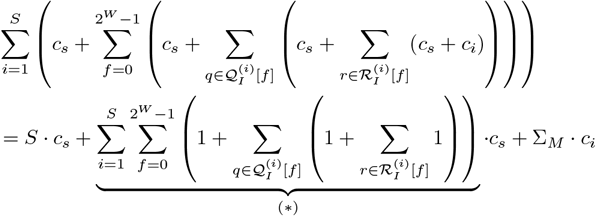

Our goal when evaluating (∗) is to avoid making the term *S*2^*W*^ appear, as it would be prohibitive; we intend to absorb this cost into the others factors. To this end, we work backward from the inner parenthesis. Let us fix 1 ≤ *i* ≤ *S*, 0 ≤ *f* ≤ 2^*W*^ − 1 and 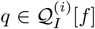 assuming it exists. We then consider the following term (*a*) 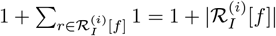. Notice that this term is equal to

- 1 if 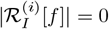,
- 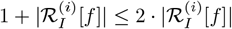 otherwise.

Therefore, (*a*) is bounded by 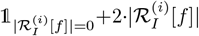. Now consider the term (*b*) :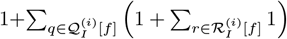.By what precedes, we have

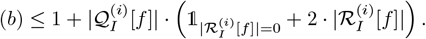

If 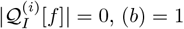. Otherwise,

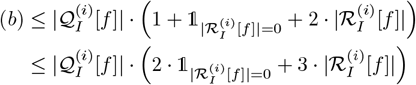

Finally, we have established 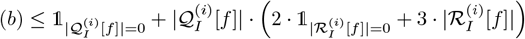 and

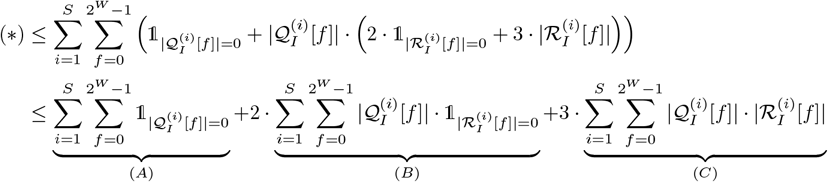

(*C*) is the easiest sum, as we use the same trick as for the proof of Proposition 3, and get (*C*) = Σ_*M*_. In Theorem 1, we made the assumption that for any document *d* ∈ 𝒟, 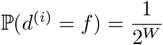. Therefore, the probability that a given fingerprint *f* is never seen in 𝒟 is 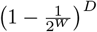, where *D* = |𝒟|. Now, using the fundamental bridge property, remark that

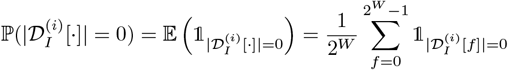

Therefore, 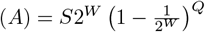. Finally, noticing that 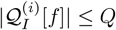 and by the same probabilistic argument, 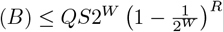.

Gathering everything together, and noticing that we could have exchanged the order among which we process 𝒬 and ℛ, we end with a complexity for Algorithm 3 of

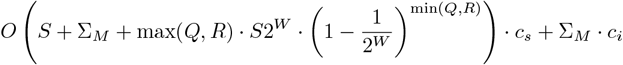

Without loss of generality, assume *Q* = *R*. Then, 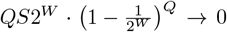 as *Q* → ∞; with the assumption that *Q* is large enough, this term is made negligible and the statement holds.

**Remark 2**. *The calculation trick that allows us to get rid of the term in S* · 2^*W*^ *in the previous proof can be interpreted as follows. We are indeed iterating over all* 2^*W*^ *possible fingerprints for each sketch. However, whenever a fingerprint f exists somewhere in some document, the cost of iteration for this particular fingerprint can be absorbed in the subsequent processing of (non-empty) list* 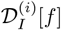. *On the other hand, if a fingerprint is not seen in any document, then we pay the price of iteration for this fingerprint, ‘for nothing’. It suffices to remark that when both Q and R are large, the probability that a fingerprint does not exist is almost zero, so that we can consider that no fingerprint is seen ‘for nothing’*.

In the end, the contribution of the sequential access cost *c*_*s*_ amounts to *O*(*QRS*) for Algorithm 1, *O*(*QS* + Σ_*M*_) for Algorithm 2, and *O*(*S* + Σ_*M*_) for Algorithm 3. Since *S* ≪ Σ_*M*_ for non-pathological cases, we end up retrieving the same comparison as is found in Table 1.

### S3 Influence of the Agievich bound on the estimation of ℙ (*X* = *k*|*p* ≥ *t*)

Figure 6 illustrates how the values 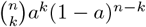 and *U* (*n, k*)*a*^*k*^(1 − *a*)^*n*−*k*^ differ.

**Figure 6:**
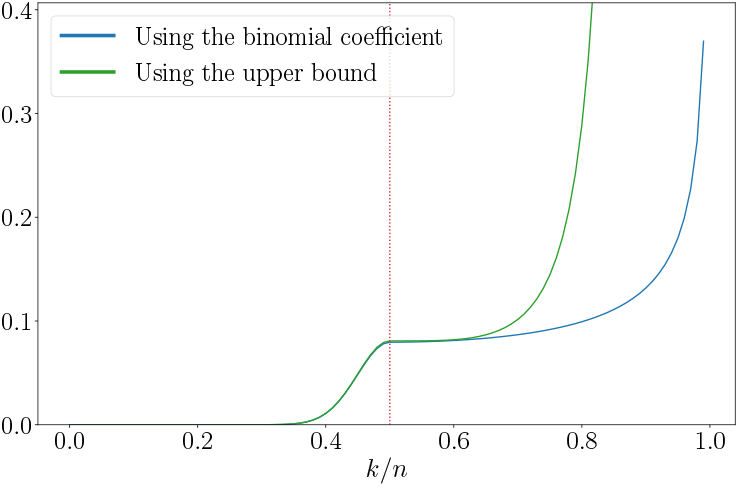
Values of the two upper bounds of ℙ (*X* = *k* | *p* ≥ *t*), with *n* = 100 and *t* = 0.5 (red vertical line).

**Figure 7:**
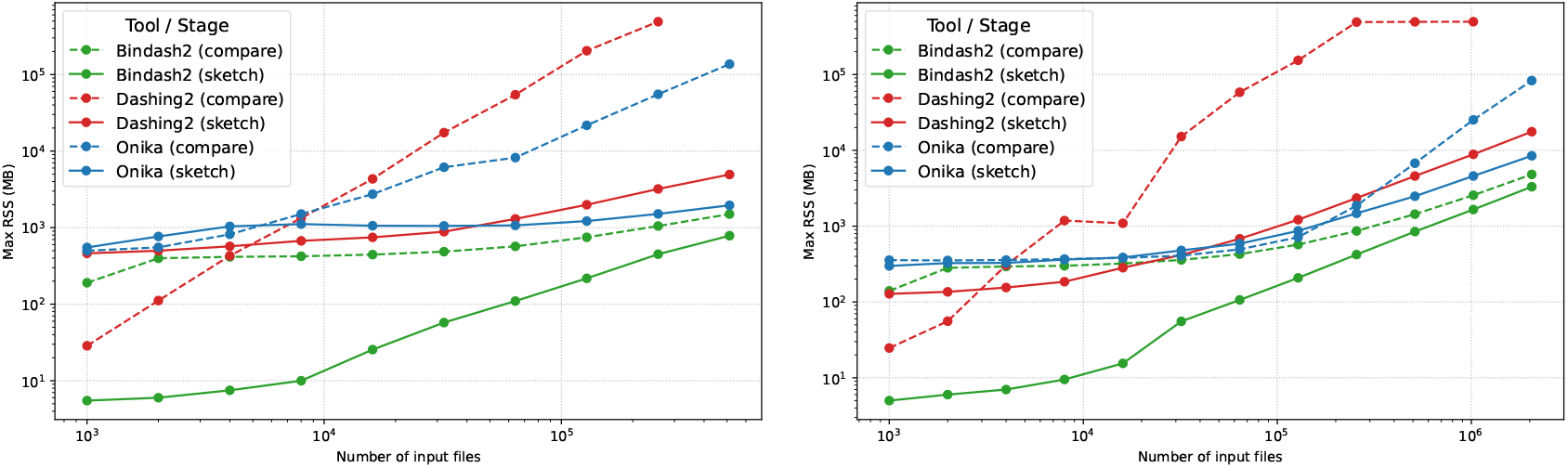
Comparison of maximum RAM usage of Dashing2, Bindash2 and Onika on a growing collection of bacterial genomes from RefSeq (left) and from random 1 Mb sequences (right). For both experiments *k* = 31 and *s* = 512

## References

[1] Chikhi, R. et al. Logan: planetary-scale genome assembly surveys life’s diversity. bioRxiv 2024–07 (2024).

[2] Karasikov, M. et al. Efficient and accurate search in petabase-scale sequence repositories. Nature 1–9 (2025).

[3] Li, S., Carss, K. J., Halldorsson, B. V., Cortes, A. & Consortium, U. B. W.-G. S. Whole-genome sequencing of half-a-million UK Biobank participants. medRxiv 2023–12 (2023).

[4] Gustafson, J. A. et al. High-coverage nanopore sequencing of samples from the 1000 Genomes Project to build a comprehensive catalog of human genetic variation. Genome research 34, 2061– 2073 (2024).

[5] Kokoris, M. et al. Sequencing by expansion (SBX)–a novel, high-throughput single-molecule sequencing technology. bioRxiv 2025–02 (2025).

[6] AB, S. T. Method and device for high throughput imaging (2021). URL https://patents.justia.com/patent/10732113. Filed August 27, 2015. Available at https://patents.justia.com/patent/10732113.

[7] AB, S. T. High throughput biochemical screening (2024). URL https://patents.justia.com/patent/10908073. Filed February 28, 2022. Available at https://patents.justia.com/patent/10908073.

[8] Zielezinski, A., Vinga, S., Almeida, J. & Karlowski, W. M. Alignment-free sequence comparison: benefits, applications, and tools. Genome biology 18, 186 (2017).

[9] Ondov, B. D. et al. Mash: fast genome and metagenome distance estimation using MinHash. Genome biology 17, 132 (2016).

[10] Shaw, J. & Yu, Y. W. Fast and robust metagenomic sequence comparison through sparse chaining with skani. Nature Methods 20, 1661–1665 (2023).

[11] Agret, C., Cazaux, B. & Limasset, A. Toward optimal fingerprint indexing for large scale genomics. In WABI 2022-22nd International Workshop on Algorithms in Bioinformatics, vol. 242, 25–1 (2022).

[12] Marchet, C. et al. Data structures based on k-mers for querying large collections of sequencing data sets. Genome research 31, 1–12 (2021).

[13] Bradley, P., Den Bakker, H. C., Rocha, E. P., McVean, G. & Iqbal, Z. Ultrafast search of all deposited bacterial and viral genomic data. Nature biotechnology 37, 152–159 (2019).

[14] Harris, R. S. & Medvedev, P. Improved representation of sequence bloom trees. Bioinformatics 36, 721–727 (2020).

[15] Marchet, C. & Limasset, A. Scalable sequence database search using partitioned aggregated Bloom comb trees. Bioinformatics 39, i252–i259 (2023).

[16] Ulrich, J.-U. & Renard, B. Y. Fast and space-efficient taxonomic classification of long reads with hierarchical interleaved XOR filters. Genome Research 34, 914–924 (2024).

[17] Marchet, C., Kerbiriou, M. & Limasset, A. BLight: efficient exact associative structure for k-mers. Bioinformatics 37, 2858–2865 (2021).

[18] Pibiri, G. E. Sparse and skew hashing of k-mers. Bioinformatics 38, i185–i194 (2022).

[19] Pibiri, G. E., Shibuya, Y. & Limasset, A. Locality-preserving minimal perfect hashing of k-mers. Bioinformatics 39, i534–i543 (2023).

[20] Broder, A. Z. On the resemblance and containment of documents. In Proceedings. Compression and Complexity of SEQUENCES 1997 (Cat. No. 97TB100171), 21–29 (IEEE, 1997).

[21] Zhao, X. BinDash, software for fast genome distance estimation on a typical personal laptop. Bioinformatics 35, 671–673 (2019).

[22] Zhao, J., Zhao, X., Pierre-Both, J. & Konstantinidis, K. T. Bindash 2.0: new MinHash scheme allows ultra-fast and accurate genome search and comparisons. bioRxiv 2024–03 (2024).

[23] Baker, D. N. & Langmead, B. Dashing: fast and accurate genomic distances with HyperLogLog. Genome biology 20, 265 (2019).

[24] Baker, D. N. & Langmead, B. Genomic sketching with multiplicities and locality-sensitive hashing using Dashing 2. Genome Research 33, 1218–1227 (2023).

[25] Ertl, O. SetSketch: filling the gap between MinHash and HyperLogLog. arXiv preprint 2101.00314 (2021).

[26] Hera, M. R., Pierce-Ward, N. T. & Koslicki, D. Deriving confidence intervals for mutation rates across a wide range of evolutionary distances using fracminhash. Genome research 33, 1061–1068 (2023).

[27] Irber, L. et al. sourmash v4: A multitool to quickly search, compare, and analyze genomic and metagenomic data sets. Journal of Open Source Software 9, 6830 (2024).

[28] Rahman Hera, M. & Koslicki, D. Estimating similarity and distance using fracminhash. Algorithms for Molecular Biology 20, 8 (2025).

[29] Rouzé, T., Martayan, I., Marchet, C. & Limasset, A. Fractional hitting sets for efficient multiset sketching. Algorithms for Molecular Biology 20, 1 (2025).

[30] Ondov, B. D. et al. Mash screen: high-throughput sequence containment estimation for genome discovery. Genome biology 20, 232 (2019).

[31] Rowe, W. P. When the levee breaks: a practical guide to sketching algorithms for processing the flood of genomic data. Genome biology 20, 199 (2019).

[32] Zobel, J. & Moffat, A. Inverted files for text search engines. In ACM Computing Surveys, vol. 38, 6 (2006). Comprehensive survey of inverted index structures, compression, and query processing techniques.

[33] Strozecki, Y. Enumeration complexity: incremental time, delay and space. arXiv preprint 2309.17042 (2023).

[34] Bayardo, R. J., Ma, Y. & Srikant, R. SScaling up all pairs similarity search. In Proceedings of the 16th international conference on World Wide Web, 131–140 (2007).

[35] Sarawagi, S. & Kirpal, A. Efficient set joins on similarity predicates. In Proceedings of the 2004 ACM SIGMOD international conference on Management of data, 743–754 (2004).

[36] Mann, W., Augsten, N. & Bouros, P. An empirical evaluation of set similarity join techniques. Proceedings of the VLDB Endowment 9, 636–647 (2016).

[37] Cohen, E. Min-hash sketches. In Encyclopedia of Algorithms, 1282–1287 (Springer, 2016).

[38] Knuth, D. E. The art of computer programming. vol. 3: Sorting and searching. Reading (1973).

[39] Janson, S., Lumbroso, J. & Sedgewick, R. Bit-array-based alternatives to HyperLogLog. In Mailler, C. & Wild, S. (eds.) 35th International Conference on Probabilistic, Combinatorial and Asymptotic Methods for the Analysis of Algorithms (AofA 2024), vol. 302 of Leibniz International Proceedings in Informatics (LIPIcs), 5:1–5:19 (Schloss Dagstuhl – Leibniz-Zentrum für Informatik, Dagstuhl, Germany, 2024). URL https://drops.dagstuhl.de/entities/document/10.4230/LIPIcs.AofA.2024.5.

[40] Li, P. & König, C. b-bit minwise hashing. In Proceedings of the 19th international conference on World wide web, 671–680 (2010).

[41] Li, P. & König, A. C. Theory and applications of b-bit minwise hashing. Communications of the ACM 54, 101–109 (2011).

[42] Mohamadi, H., Chu, J., Vandervalk, B. P. & Birol, I. ntHash: recursive nucleotide hashing. Bioinformatics 32, 3492–3494 (2016).

[43] Kazemi, P. et al. ntHash2: recursive spaced seed hashing for nucleotide sequences. Bioinformatics 38, 4812–4813 (2022).

[44] Vandamme, L., Cazaux, B. & Limasset, A. K2R: Tinted de bruijn graphs implementation for efficient read extraction from sequencing datasets. Bioinformatics Advances 5, vbaf111 (2025).

[45] Girard, M., Vandamme, L., Cazaux, B. & Limasset, A. OReO: optimizing read order for practical compression. Bioinformatics Advances 5, vbaf128 (2025).

[46] Agievich, S. V. An upper bound on binomial coefficients in the de Moivre Laplace form. Journal of the Belarusian State University. Mathematics and Informatics 1, 66–74 (2022).

[47] Berlin, K. et al. Assembling large genomes with single-molecule sequencing and locality-sensitive hashing. Nature biotechnology 33, 623–630 (2015).

[48] Shafin, K. et al. Nanopore sequencing and the Shasta toolkit enable efficient de novo assembly of eleven human genomes. Nature Biotechnology 38, 1044–1053 (2020). URL https://www.nature.com/articles/s41587-020-0503-6.

[49] Koren, S. et al. Canu: scalable and accurate long-read assembly via adaptive k-mer weighting and repeat separation. Genome research 27, 722–736 (2017).

[50] Clementi, A., Gualà, L., Pepè Sciarria, L. & Straziota, A. Maintaining k-MinHash signatures over fully-dynamic data streams with recovery. In Proceedings of the Eighteenth ACM International Conference on Web Search and Data Mining, 79–87 (2025).

[51] Son, Y., Kim, C. & Lee, J. FED: Fast and efficient dataset deduplication framework with GPU acceleration (2025). URL https://arxiv.org/abs/2501.01046. ArXiv preprint, posted 2025-01-02, 2501.01046.

[52] Wald, A. On cumulative sums of random variables. The Annals of Mathematical Statistics 15, 283–296 (1944).

[53] Turner, A. L., Uppal, A. & Xu, P. Spacing distribution of a bernoulli sampled sequence. arXiv preprint 1510.03500 (2015).

